# The Global Virome in One Network (VIRION): an atlas of vertebrate-virus associations

**DOI:** 10.1101/2021.08.06.455442

**Authors:** Colin J. Carlson, Rory J. Gibb, Gregory F. Albery, Liam Brierley, Ryan P. Connor, Tad A. Dallas, Evan A. Eskew, Anna C. Fagre, Maxwell J. Farrell, Hannah K. Frank, Renata L. Muylaert, Timothée Poisot, Angela L. Rasmussen, Sadie J. Ryan, Stephanie N. Seifert

## Abstract

Data cataloguing viral diversity on Earth have been fragmented across sources, disciplines, formats, and various degrees of open collation, posing challenges for research on macroecology, evolution, and public health. Here, we solve this problem by establishing a dynamically-maintained database of vertebrate-virus associations, called The Global Virome in One Network (VIRION). The VIRION database has been assembled through both reconciliation of static datasets and integration of dynamically-updated databases. These data sources are all harmonized against one taxonomic backbone, including metadata on host and virus taxonomic validity and higher classification; additional metadata on sampling methodology and evidence strength are also available in a harmonized format. In total, the VIRION database is the largest open-source, open-access database of its kind, with roughly half a million unique records that include 9,521 resolved virus “species” (of which 1,661 are ICTV ratified), 3,692 resolved vertebrate host species, and 23,147 unique interactions between taxonomically-valid organisms. Together, these data cover roughly a quarter of mammal diversity, a tenth of bird diversity, and ~6% of the estimated total diversity of vertebrates, and a much larger proportion of their virome than any previous database. We show how these data can be used to test hypotheses about microbiology, ecology, and evolution, and make suggestions for best practices that address the unique mix of evidence that coexists in these data.

## Introduction

The global virome — the aggregate of all viruses across the entire biosphere — is one of the most under-documented components of global biodiversity. There are at least 40,000 species of viruses estimated to infect mammals alone, of which thousands can probably infect humans (Carlson et al., 2019); thousands of others, if not millions, are distributed across the tree of life, of which a select few can cross between branches with a deep evolutionary split. For example, both influenza viruses and coronaviruses circulate among birds, terrestrial mammals, and cetaceans (Krammer et al., 2018); similarly, West Nile virus infects not only humans and other mammals but also birds (Marra et al., 2003) and even some reptiles and amphibians (Klenk and Komar, 2003). In this regard, the global virome forms a broad, tangled network of species interactions — a network that is substantially under-documented.

To date, most data sources that describe the animal-virus network have been limited to a small proportion of known mammal viruses, covering at most 1-2% of the total viral diversity in this class; data on the other vertebrate classes are even more limited. These data are a critical piece of zoonotic risk assessment, as viruses capable of broad host jumps are those most predisposed to future emergence. Moreover, recent evidence shows that data on the network structure of the global virome can be used to improve predictions of viruses with zoonotic potential based on genome composition (Poisot et al., 2021). Despite the obvious value of host-virus network data, currently available datasets are notably limited by the challenges of data integration and reconciliation; manual re-compilation and reconciliation is too inefficient to consistently keep these datasets up to date, and many sources are effectively a decade behind current knowledge (Gibb et al., 2021). Therefore, open, reproducible data that are as complete and detailed as possible are desperately needed.

Here, we describe a new open database called The Global Virome in One Network (VIRION), which integrates different existing datasets to produce the most comprehensive picture ever developed of the global vertebrate virome. A total of seven data sources on host-virus associations (evidence that a virus is capable of infecting, or is known to infect, a given host) are integrated in a pipeline that standardizes taxonomy and harmonizes sampling metadata (Figure 1). The current version of the data product, and disaggregated datasets for more informal inquiry, are all openly available with this publication.

**Figure 1.**
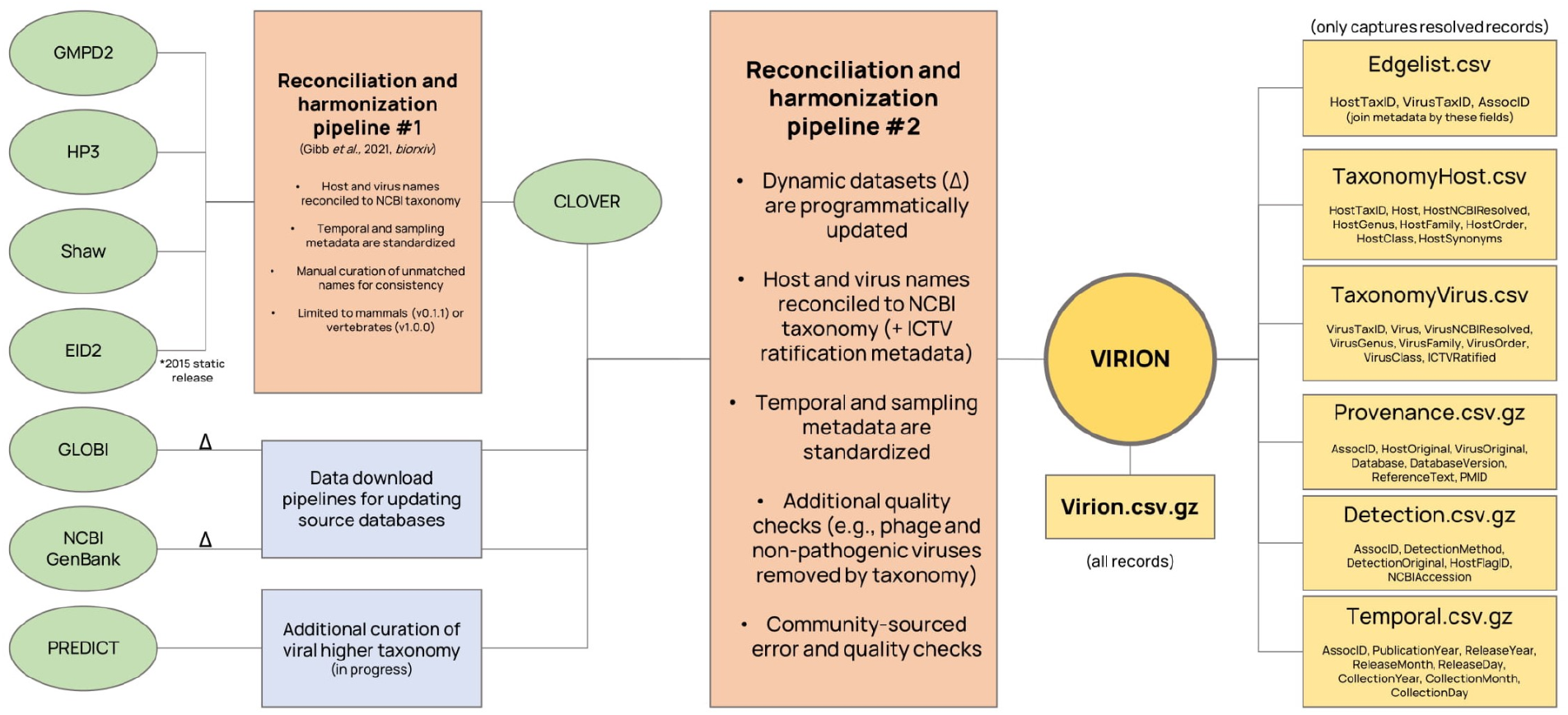
The VIRION pipeline. Data is integrated from a total of seven sources into one master file and a set of six disaggregated files that constitute the VIRION database. Data sources marked with a delta can be dynamically updated as new data are submitted to the source databases.

## Data Sources and Tailored Curation

VIRION aggregates information from a total of five static sources (referred to as HP3, GMPD2, EID2, Shaw, and PREDICT) and two dynamic sources (Global Biotic Interactions and GenBank, hosted by the National Center for Biotechnology Information [NCBI]). The first four of these sources are separately reconciled in a new version of the CLOVER database (Gibb et al., 2021), which previously only covered mammalian viruses, but is here expanded to all vertebrate hosts; the rest of the data sources are integrated into the same data architecture. There is a high degree of redundancy among these different sources, both because they draw on the same limited body of scientific literature, and because they may directly query or incorporate each other (e.g., many of the viral genome sequences generated by the PREDICT program have been deposited in GenBank). However, there are also significant differences in coverage between these data sources, and their reconciliation can reduce data sparsity by 30% or more (Gibb et al., 2021).

### The expanded CLOVER database

In a previous study (Gibb et al., 2021), we presented a semi-manually curated dataset called CLOVER (version 0.1.0) that reconciled four sources of information on mammal-virus interactions: the Host-Pathogen Phylogeny Project (HP3) database (Olival et al., 2017), the Global Mammal Parasite Database 2.0 (Stephens et al., 2017), the static version of the ENHanCEd Infectious Disease Database (EID2) from its original open release (Wardeh et al., 2015), and an unnamed dataset developed by Shaw *et al.* referred to by its lead author (Shaw et al., 2020). Each of these datasets has been widely used for hypothesis testing in viral macroecology and zoonotic risk assessment. In the updated CLOVER dataset (version 1.0.0) that we introduce here, the purview of this reconciliation is expanded to all vertebrate hosts available in the four databases (two of which, EID2 and Shaw, include non-mammalian hosts). As with CLOVER, three of these databases (HP3, GMPD2, and Shaw) are static sources, while EID2 is dynamically updated but cannot be downloaded in a comprehensive format, limiting our efforts to the 2015 static release (Wardeh et al., 2015).

The reconciliation process was the same as described in the original CLOVER publication (Gibb et al., 2021). First, virus names were manually standardized across all four datasets to ensure consistency in naming convention. Then we used the R package ‘taxize’ (Chamberlain and Szöcs, 2013) to automatically match all host and virus names to the NCBI taxonomic backbone and access higher taxonomy. Remaining host and virus names without an exact match were then manually reconciled. 99.3% of hosts (n = 2,327) and 97.7% of viruses (n = 973) were matched to a taxonomic identifier in the NCBI database; higher taxonomy for the remaining unmatched 16 hosts and 23 viruses were manually curated by a comparison against other data sources (the IUCN Red List database and a mammal phylogenetic supertree (Upham et al., 2019)). Virus detection methods as reported for each association were standardised to a 4-tier classification system: either 1) antibodies (i.e., serological evidence), 2) genetic sequencing (PCR or related methods), 3) viral isolation, or 4) not reported. Temporal metadata (year of accession or publication) were accessed for each association, either from the reported primary source publication, or by querying associated PubMed and NCBI Nucleotide records using the R package ‘rentrez’ (Winter, 2017). The reconciled CLOVER dataset contained a total of 70,466 records comprising 8,004 unique host-virus associations.

### The USAID PREDICT testing database

The USAID Emerging Pandemic Threats (EPT) PREDICT program ran from 2009 to 2019, and remains the largest coordinated program of wildlife viral discovery to date, with a roughly $200m USD investment that led to the identification of 218 known and 931 novel viruses (Carlson, 2020; Grange et al., 2021). In 2021, data generated by this program were released on the USAID data library and an API through HealthMap (healthmap.org/predict), which offer largely overlapping versions of the same data. Many, but not all, of these sequences have been uploaded to GenBank (see below), while others are already present in literature-based databases, but a number of novel records are also given in these newly-released data.

We obtained all the data available through the HealthMap public API, which includes a slightly greater coverage of data than the USAID data library release. We then downloaded the USAID version of the data library once released, and added any associations not present in the other dataset. All host names were already cleaned and formatted for reuse, though a handful of records were manually flagged for uncertain host identification; in PREDICT studies, some hosts were identified solely by relying on morphological traits observed in the field, while others had identification confirmed via genetic barcoding. Some of these are denoted by an asterisk in the HealthMap data copy, while others are given as “Genus cf. species” in both data copies; in both cases, we retained this self-reported information in the ‘HostFlagID’ field as a measure of uncertainty.

Virus names required a more detailed reconciliation workflow. First, virus names were simplified (i.e., influenza sequence details or similar lineage information were dropped). Subsequently, we validated virus names first by querying the NCBI taxonomy directly (using the ‘taxize’ R package), and subsequently by retrieving the metadata for GenBank accessions that are reported in the PREDICT dataset (using the ‘rentrez’ R package). A number of names still remain unresolved, including those that follow the bespoke naming convention used by the PREDICT program. These names are assigned the lowest possible rank of higher taxonomy based on their naming convention; for example, PREDICT_MAstV-161 is assigned to the genus *Mamastrovirus* (Stellavirales: Astroviridae), while PREDICT_CoV-24 is only assigned to the family Coronaviridae (Nidovirales). For many of the viruses discovered by the PREDICT project, additional genus information is available from the SpillOver Risk Ranking Tool database (spillover.global; (Grange et al., 2021)). We downloaded these data and matched them to the names in the PREDICT data releases where possible. Future efforts may be able to improve taxonomic resolution for the remaining viruses based on phylogenetic analysis using currently-unreleased sequence data; these will be incorporated through GenBank accessions, rather than updates to the PREDICT datasets, which are currently presumed to be static releases.

### Global Biotic Interactions (GLOBI)

GLOBI is a dynamic, content-aggregating, open-access database of ecological interactions(Poelen et al., 2014). The data in GLOBI are automatically pulled and manually curated from published scientific literature and other databases, including the bat virus database DBatVir (www.mgc.ac.cn/DBatVir), the rodent virus database DRodVir (www.mgc.ac.cn/DRodVir), and Virus-Host DB (www.genome.jp/virushostdb). As such, GLOBI collates a number of data sources with value for describing the global virome. However, in GLOBI, each species’ taxonomic identity is linked to the initial taxonomic database to which it was reconciled, representing a number of different backbones (e.g., NCBI, GBIF, Encyclopedia of Life) each of which coexists in these datasets. As such, these data may include duplications, outdated synonyms, or other inconsistencies: for example, “Virus” and “Viruses” are both used as high-level ranks.

We use the R package ‘rglobi’ (Poelen et al., 2021) to automatically download all associations between viruses (queried as both “Virus” and “Viruses”) and vertebrates (queried as “Vertebrata”) in the dataset. Virus names are simplified (i.e., influenza A/B/C/D sequence details or similar lineage information are dropped), and both host and virus names are validated against the NCBI taxonomic backbone. In an effort to standardize data quality, we only retain hosts that can be matched (and confirmed as vertebrates) against the NCBI backbone.

### NCBI GenBank

GenBank is the gold-standard, open-access, community-generated dataset of nucleotide and protein sequence data for life on Earth, with over two billion sequences submitted as of February 2021. For viruses, there are nearly five million distinct accessions of nucleotide sequences, which can be directly downloaded from the NCBI Virus interface (https://www.ncbi.nlm.nih.gov/labs/virus) or from an FTP server (https://ftp.ncbi.nlm.nih.gov/genomes/Viruses/AllNuclMetadata/). Though the majority of viral samples (more than 11 million) are from *Homo sapiens*, and many others are from bacteriophages, GenBank is still an incomparable source of information on vertebrate-virus associations, with genetic sequence representing strong evidence in support of these associations.

For the VIRION workflow, we download the entirety of NCBI Virus’s nucleotide records. While virus samples in GenBank are almost all resolved to the NCBI taxonomic backbone, host information is contributed by users directly as sample metadata, and is less standardized. We therefore used the R package ‘taxize’ to clean host metadata and obtain higher classification data for both hosts and viruses. To ensure data quality, we limited the records incorporated into VIRION to those that had a confirmed match to a vertebrate host.

### Data integration and taxonomic reconciliation

We developed a reproducible, open pipeline to integrate these seven data sources in the statistical software ‘R’. Throughout, we used a standardized set of tools and procedures, relying heavily on a set of tools for biological data science developed by the ROpenSci program.

The primary component of the data is a list of hosts and viruses known to associate with each other. Where possible, these are resolved down to the species level for both (in the fields “Host” and “Virus”); otherwise, information is given down to the lowest possible valid taxonomic rank (e.g., *Betacoronavirus* in “VirusGenus” or Chiroptera in “HostClass”), and original entries are retained (“HostOriginal”, “VirusOriginal”). All of these fields are stored in all-lowercase letters to avoid inconsistencies in capitalization. Information is also recorded on whether species names are considered valid in the NCBI and International Committee on the Taxonomy of Viruses (ICTV) taxonomies, and whether host identification was originally reported as uncertain.

Unlike other data sources that reproduce taxonomic information verbatim, or use different taxonomic backbones on a per-data-point basis (e.g., GLOBI), VIRION is fully reconciled against the taxonomic hierarchy curated by the NCBI for both animals and viruses, using the R package ‘taxize’ (Chamberlain and Szöcs, 2013) to access the NCBI API. For animal hosts, the classification of major lineages generally follows the broad consensus of names selected under the appropriate code of nomenclature in recent systematic studies, while taking a conservative approach to areas that remain contentious or unresolved. For viruses, the taxonomic hierarchy is dynamically updated (and these updates propagated) to be fully harmonized with the contemporary decisions of the ICTV; although the parsing and mapping of new names to the taxonomic backbone are done automatically, the names are only propagated after review by the NCBI Taxonomy group. In addition, users can raise issues of taxonomic mismatch between NCBI and ICTV directly with the NCBI Taxonomy group, which allows these issues to be solved based on community feedback when supported by an authority; this ensures that issues identified by the community do not overrule ICTV or ICZN.

In addition to taxonomic information, a minimum standard of metadata is retained from all sources (Table 1), and integrated in a data harmonization pipeline modeled off of the CLOVER pipeline. This includes information on the source of the data; the date of sample collection, data release, or related scientific publication; detection methods; and other citation-related or accession-related information.

**Table 1.**
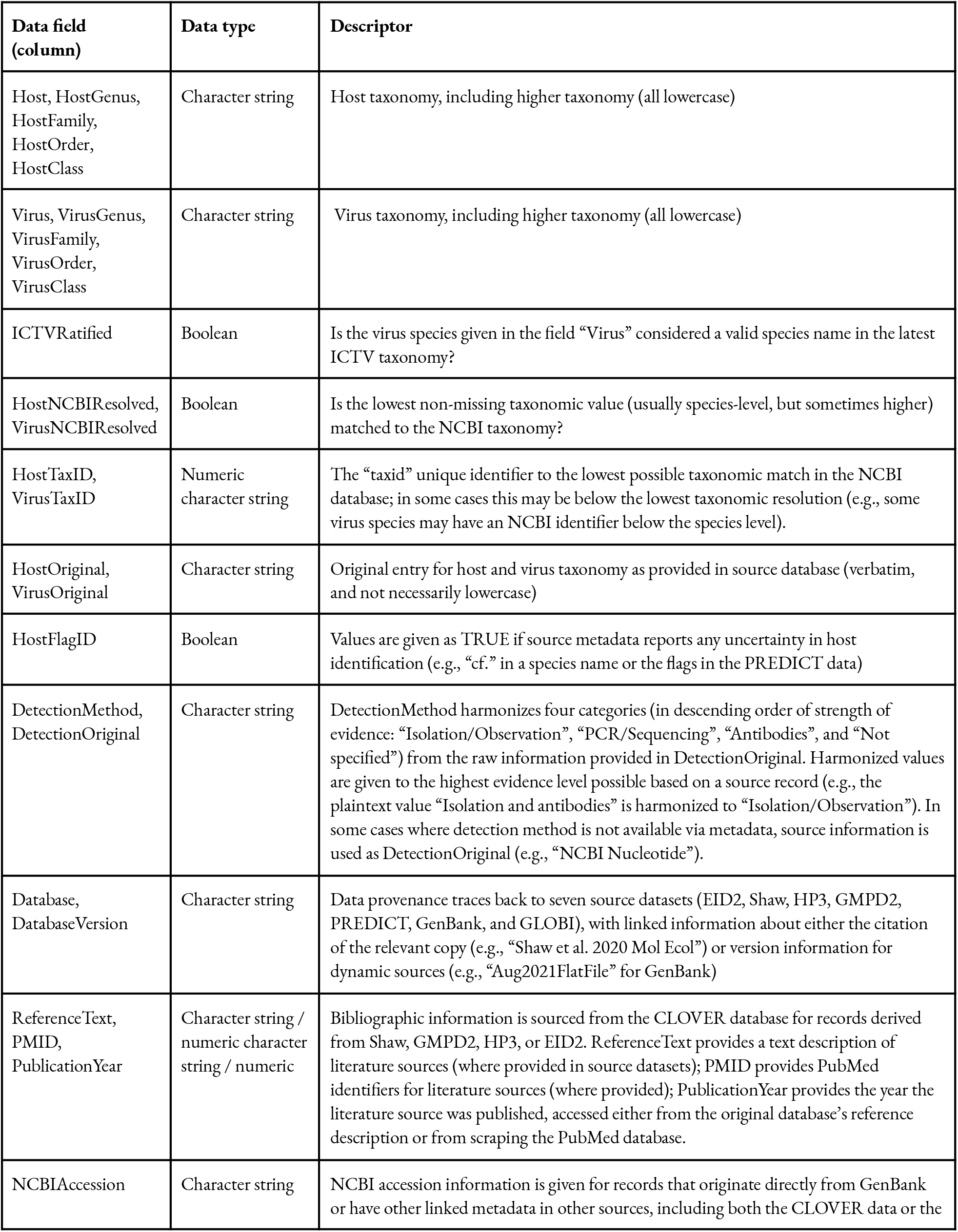
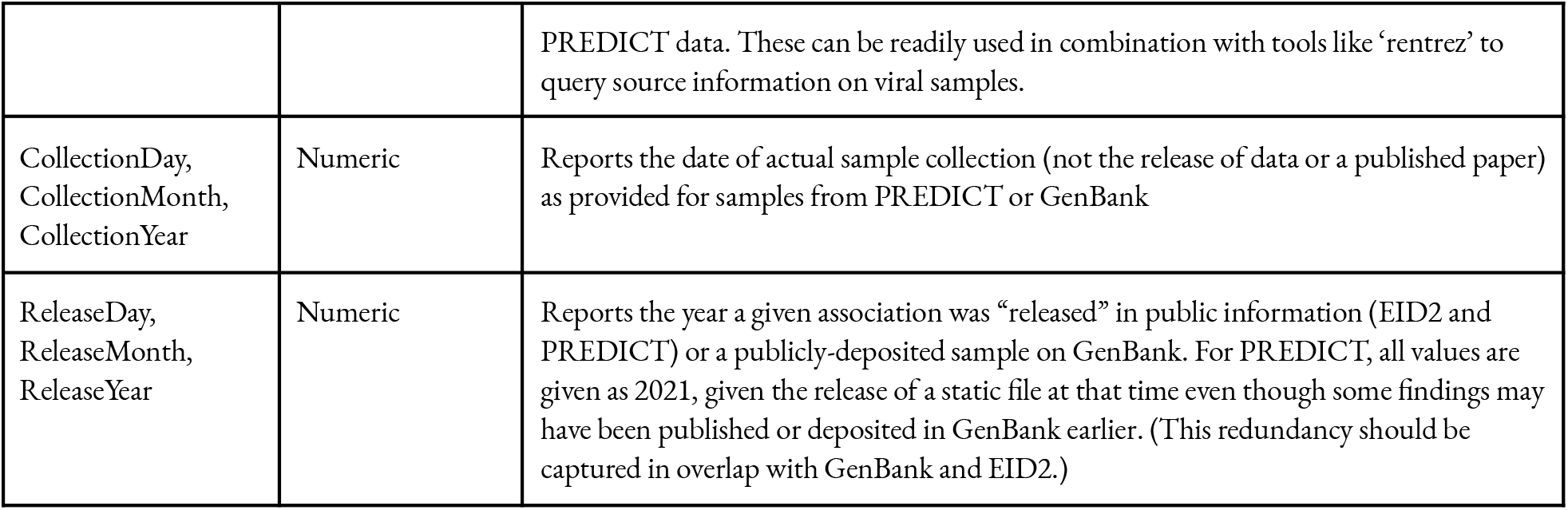
Data field descriptors for the VIRION database.

| Data field (column) | Data type | Descriptor |
| --- | --- | --- |
| Host, HostGenus, HostFamily, HostOrder, HostClass | Character string | Host taxonomy, including higher taxonomy (all lowercase) |
| Virus, VirusGenus, VirusFamily, VirusOrder, VirusClass | Character string | Virus taxonomy, including higher taxonomy (all lowercase) |
| ICTVRatified | Boolean | Is the virus species given in the field “Virus” considered a valid species name in the latest ICTV taxonomy? |
| HostNCBIResolved, VirusNCBIResolved | Boolean | Is the lowest non-missing taxonomic value (usually species-level, but sometimes higher) matched to the NCBI taxonomy? |
| HostTaxID, VirusTaxID | Numeric character string | The “taxid” unique identifier to the lowest possible taxonomic match in the NCBI database; in some cases this may be below the lowest taxonomic resolution (e.g., some virus species may have an NCBI identifier below the species level). |
| HostOriginal, VirusOriginal | Character string | Original entry for host and virus taxonomy as provided in source database (verbatim, and not necessarily lowercase) |
| HostFlagID | Boolean | Values are given as TRUE if source metadata reports any uncertainty in host identification (e.g., “cf.” in a species name or the flags in the PREDICT data) |
| DetectionMethod, DetectionOriginal | Character string | DetectionMethod harmonizes four categories (in descending order of strength of evidence: “Isolation/Observation”, “PCR/Sequencing”, “Antibodies”, and “Not specified”) from the raw information provided in DetectionOriginal. Harmonized values are given to the highest evidence level possible based on a source record (e.g., the plaintext value “Isolation and antibodies” is harmonized to “Isolation/Observation”). In some cases where detection method is not available via metadata, source information is used as DetectionOriginal (e.g., “NCBI Nucleotide”). |
| Database, DatabaseVersion | Character string | Data provenance traces back to seven source datasets (EID2, Shaw, HP3, GMPD2, PREDICT, GenBank, and GLOBI), with linked information about either the citation of the relevant copy (e.g., “Shaw et al. 2020 Mol Ecol”) or version information for dynamic sources (e.g., “Aug2021FlatFile” for GenBank) |
| ReferenceText, PMID, PublicationYear | Character string / numeric character string / numeric | Bibliographic information is sourced from the CLOVER database for records derived from Shaw, GMPD2, HP3, or EID2. ReferenceText provides a text description of literature sources (where provided in source datasets); PMID provides PubMed identifiers for literature sources (where provided); PublicationYear provides the year the literature source was published, accessed either from the original database’s reference description or from scraping the PubMed database. |
| NCBIAccession | Character string | NCBI accession information is given for records that originate directly from GenBank or have other linked metadata in other sources, including both the CLOVER data or the |
|  |  | PREDICT data. These can be readily used in combination with tools like ‘rentrez’ to query source information on viral samples. |
| CollectionDay,<br>CollectionMonth,<br>CollectionYear | Numeric | Reports the date of actual sample collection (not the release of data or a published paper) as provided for samples from PREDICT or GenBank |
| ReleaseDay,<br>ReleaseMonth,<br>ReleaseYear | Numeric | Reports the year a given association was “released” in public information (EID2 and PREDICT) or a publicly-deposited sample on GenBank. For PREDICT, all values are given as 2021, given the release of a static file at that time even though some findings may have been published or deposited in GenBank earlier. (This redundancy should be captured in overlap with GenBank and EID2.) |

## The assembled VIRION database

VIRION is the most comprehensive open-access database characterizing the vertebrate virome. Version 0.2.1 of the database, which is released alongside the initial preprint, contains a total of 484,464 records. The majority of distinct records (n = 391,487; ~81%) come from GenBank, particularly due to human samples with distinct dates of a handful of very well sampled viruses (e.g., Influenza A virus, HIV-1, SARS-CoV-2, and Norfolk virus). The remaining 20% or so of the data are distributed unevenly among EID2 (n = 57,265; 12%), GLOBI (19,661; 4%), Shaw (8,643; 2%), PREDICT (2,850; < 1%), HP3 (2,802; <1%), and GMPD2 (1,765; < 1%). After removing metadata, distinct values of resolved host-virus associations are more evenly distributed among GenBank (15,467; 36%), GLOBI (13,924; 33%), Shaw (6,418; 15%), HP3 (2,759; 6%), EID2 (2,025; 5%), PREDICT (2%), and GMPD2 (2%).

The VIRION database is far more extensive than many of these source datasets on their own, and is information-dense with respect to associations and not just species (Figure 2). The association data include 9,521 resolved virus “species” (of which 1,661 are ICTV ratified), 3,692 resolved vertebrate host species, and 23,147 unique interactions between taxonomically-valid organisms. Together, these data cover ~25% (1,635 spp.) of mammal diversity (~6,500 spp.); ~11% (1,072 spp.) of birds (~10,000 spp.); and ~6% of the estimated total diversity of vertebrates (~60,000 spp.). Each of these groups has its own unique coverage biases and gaps in terms of both host taxonomy (Figure 3) and geographic sampling (Figure 4), which are non-trivial enough to shape the findings of research that uses the database.

**Figure 2.**
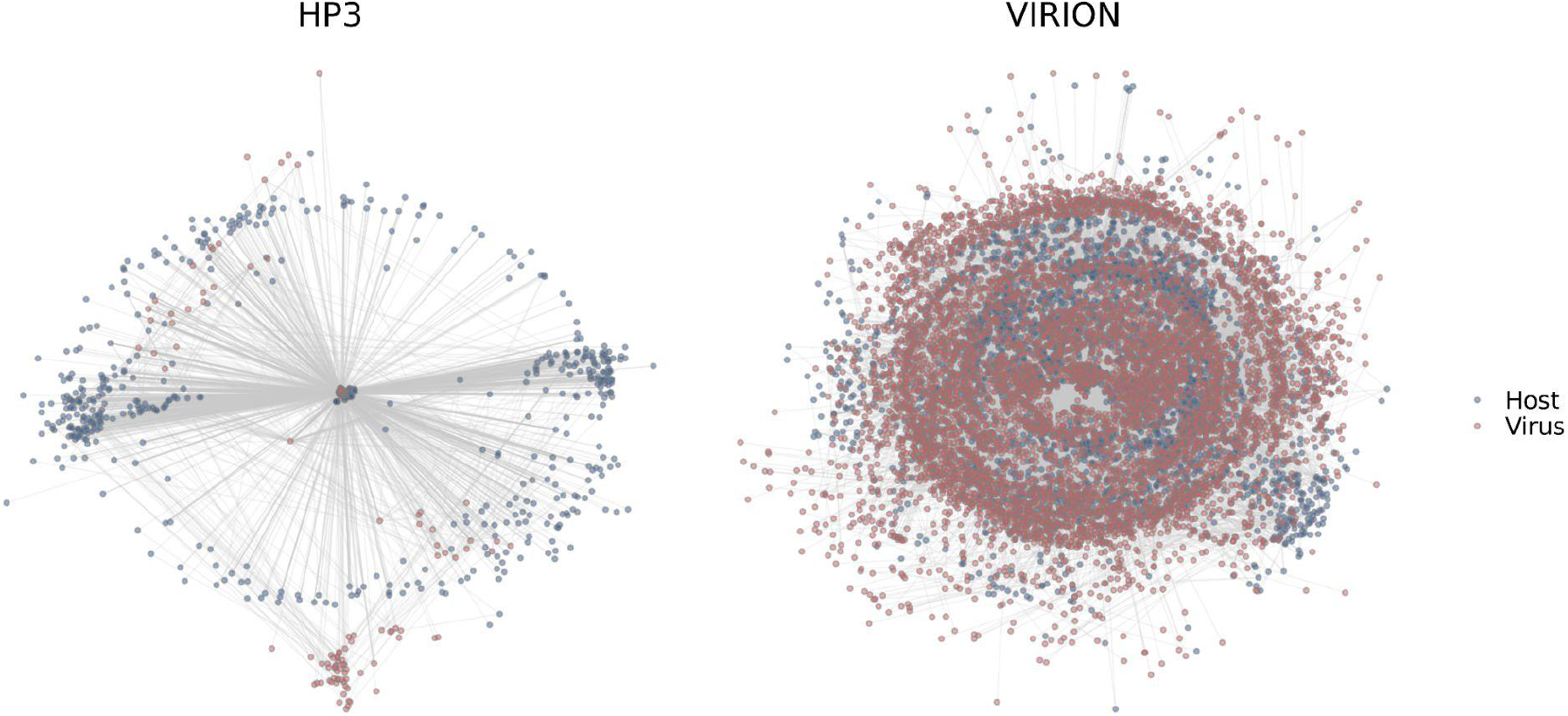
Comparative scope of data. Networks show all unique NCBI-recognized host-virus species pairs (viruses are red, hosts are blue) in the Host-Pathogen Phylogeny Project database (HP3, published in 2017) and VIRION. The information stored in VIRION is more extensive (including all vertebrates, not just mammals) but also far more information-dense, describing a network with many more nodes and many more connections.

**Figure 3.**
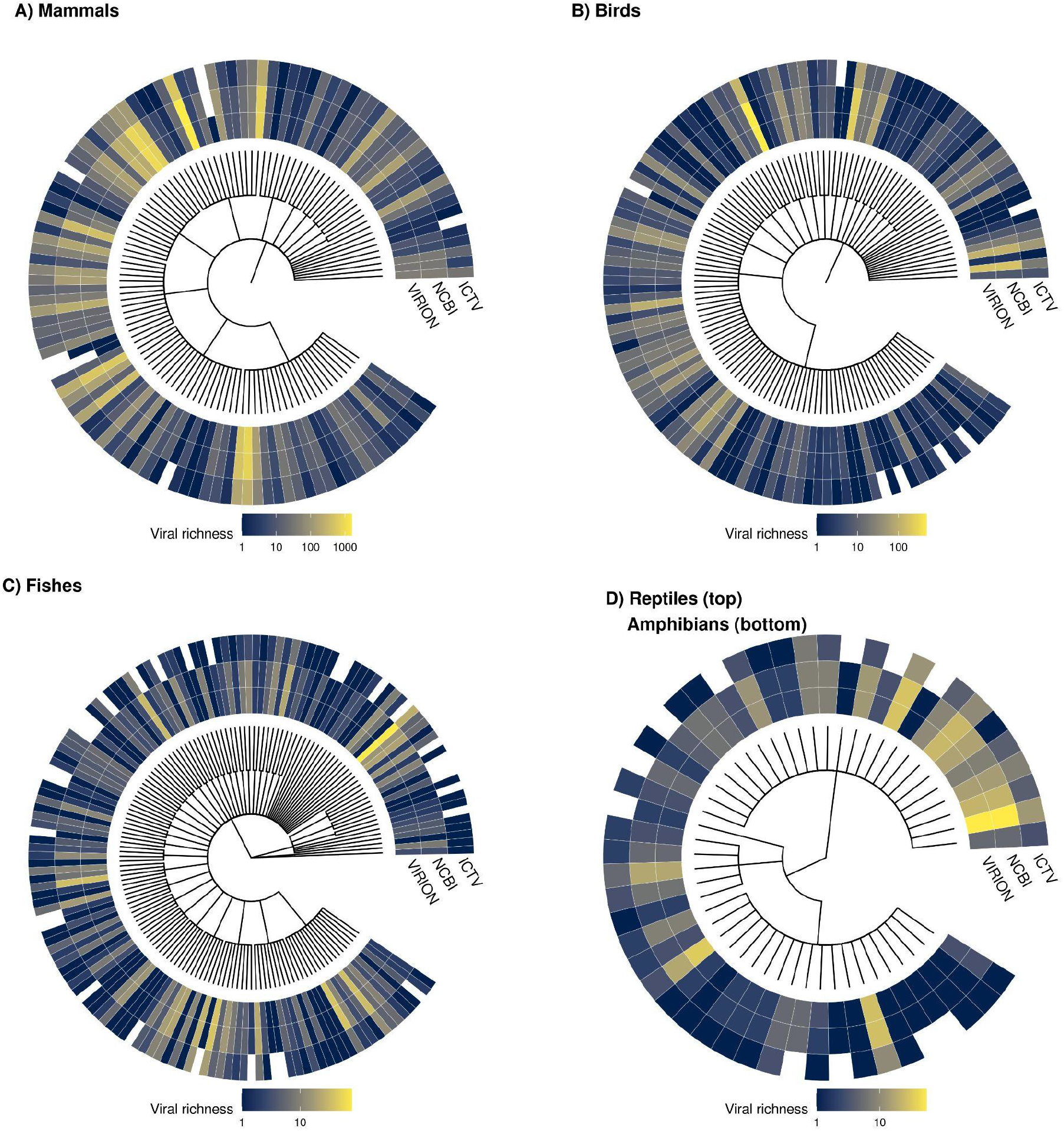
Taxonomic coverage across hosts. Each tree tip represents one host family, with the total number of viruses recorded in VIRION, the number that are NCBI-resolved, and the number that are ICTV ratified. Note that the color scale varies across panels.

**Figure 4.**
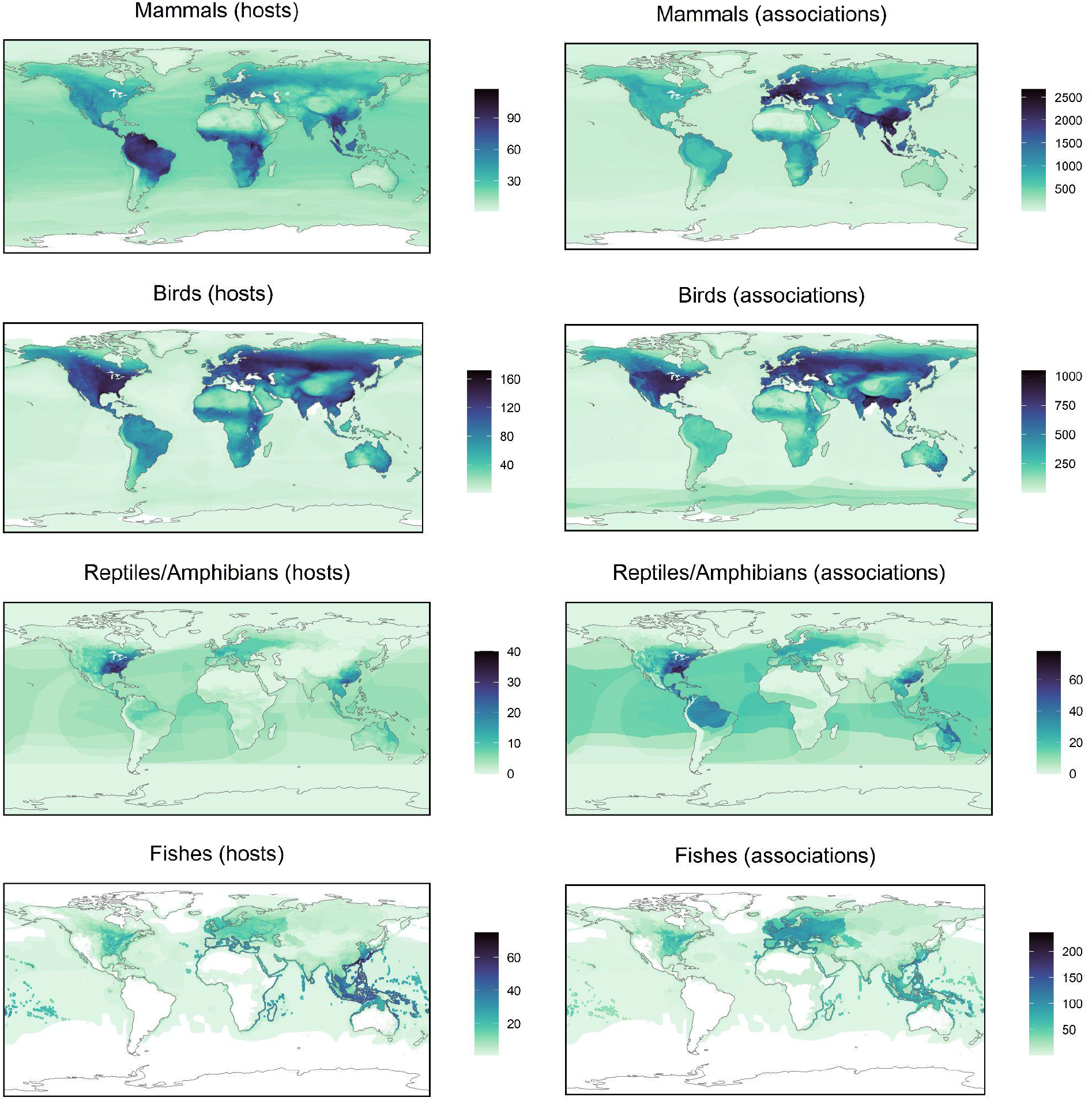
The geographic distribution of hosts andhost-virus associations based on IUCN geographic range maps. Species are matched to the IUCN database using verbatim Latin names, without any manual correction. This is largely congruent for mammals (91.1%) and birds (93%) but less so for reptiles/amphibians (79%) and fish (47.6%), in part because some species may not yet be mapped. Particularly when working with the latter groups, users will likely need to manually *cross-reference species names from the VIRION database to other sources.*

## How to use VIRION

VIRION can be downloaded from an open-source, open-access, version-labelled GitHub repository, accessible directly from the URL (github.com/viralemergence/virion) or through a link on the Verena Consortium website (viralemergence.org/virion). While the database can be used fairly out-of-the-box thanks to the taxonomic reconciliation pipeline, we encourage users to think carefully about the data they are manipulating, and how it relates to their analysis. Here, we recommend some simple steps towards that end, and show some example uses (based on release v0.2.1).

### Some best practices

We recommend that users should consider the following steps as part of their workflow:

1. **Be aware of the challenges of big data.** Make sure that column types are read in correctly if columns are particularly sparse, be aware of the challenges of reading and writing large files (packages like ‘data.table’ and ‘vroom’ in R; or ‘DataFrames.jl’, ‘JuliaDB.jl’, and ‘Arrow.jl’ in Julia; help with this), and note the small handful of optimizations we’ve made to make VIRION easier to work with (in particular, NCBIAccession values are stored as comma-space separated lists - e.g., “KP272011.1, KP272012.1” - for records that are otherwise identical in all other metadata).
2. **Consider taxonomic resolution.** Most users will only want to use records where both the virus and host names are NCBI resolved; though these are relatively standardized, many still include sampling metadata or unclassified lineage information for viruses. Simple rubrics can flag these cases: for example, in the list of NCBI-accepted betacoronavirus names, searching for virus names that include a “/” will flag many lineage-specific records (“bat coronavirus 2265/philippines/2010”, “coronavirus n.noc/vm199/2007/nld“) and separate them from some cleaner names (“alpaca coronavirus“), but won’t necessarily catch everything (“bat coronavirus ank045f“). Another option is to limit analysis to viruses that are ICTV ratified (“ICTVRatified” = TRUE), but this is particularly conservative, and will leave a much larger number of valid virus names out (Fig. 3). Users should consider a read through of all of the species names in the dataset they’re working with, just to familiarize themself with any remaining oddities. (When errors or oddities are detected in the NCBI taxonomy, they can be submitted for investigation and a possible patch on the GitHub repository; these issues should be tagged using the label “taxonomy”.)
3. **Consider paraphyletic relationships.** The NCBI taxonomic backbone uses no-rank “in part” names to denote paraphyletic groups; these ranks may not be found consistently at higher taxonomic levels that belong to the paraphyletic group. In a situation where a user would query all viruses for a paraphyletic group like Reptilia, this can lead to missing hosts. An alternative more robust practice would be to query the component taxa directly, *i.e.* Lepidosauria (rank: class; TaxID: 8504) and Testudines (order; 8459) and Crocodylia (order; 1294634). These issues are likely to be more prevalent at high taxonomic ranks in the host taxonomy.
4. **Consider data origins and sampling.** The datasets in CLOVER contain rich information on which associations are known from serology, sequencing/PCR, or viral isolation. Other data sources contain less rich information: GLOBI does not retain this sampling metadata, and all GenBank records are assigned to sequencing as the “minimum evidence standard” even though some may be viral isolation. In combination with other nuances about data provenance, this can also indicate differences in the underlying biological reality: for example, HP3, Shaw, and GMPD2 are manually curated to only include data from wild, “natural” infections, and explicitly exclude experimental infections; EID2, which automatically pulls in new records from sources like GenBank, contains some experimental infections (e.g., baboons and macaques in the United States). A researcher interested in host-virus biological compatibility may be fine using any of the data sources in VIRION, while one trying to document patterns in the wild may wish to subset based on detection and data provenance accordingly.
5. **Check for quality issues programmatically.** For example, “HostFlagID” denotes the presence of possible uncertainty in host identification, which users may want to check before proceeding any further. This uncertainty may or may not be acceptable depending on data provenance and user intentions.
6. **Check for quality issues manually when possible.** For the most important applications, narrowly defining the dataset of interest can help users make the task of manual quality control a manageable one. For example, if a researcher is developing a predictive model of the host-virus network for the purpose of identifying viruses with zoonotic potential, they may want to manually check each virus that is recorded as having human hosts (see Example 2), which will be the most important subset of the data for model performance.

Finally, we encourage users who discover unusual interactions to investigate their provenance and potentially flag controversial data in source datasets; this can be particularly important for the most valuable records in VIRION, such as those that document zoonotic infections. Erroneous records of human infection are likely to significantly impact models and may even lead to public health confusion, and as such, data should be carefully handled and reproduced in analytical settings.

#### Example 1: Where does chikungunya come from?

In the early days of the Verena Consortium (viralemergence.org), a debate started in the margins of a heavily-edited manuscript: is it appropriate to include chikungunya in a list of zoonotic diseases? Chikungunya virus is a mosquito-borne alphavirus of particular concern to public health in the tropics; unlike many comparable alphaviruses (e.g., Mayaro virus) and flaviviruses (e.g., Zika virus), chikungunya is rarely thought of as a pathogen with a sylvatic cycle in non-human primates. But like most *Aedes-*borne viruses, it can certainly infect some animals, having been isolated from non-human primates (Apandi et al., 2009), bats (Apandi et al., 2009; Zhang et al., 1989), and birds (Moore et al., 1974) in the wild. Further, experimental challenge studies indicate susceptibility of additional bat species, amphibians, and reptiles to chikungunya virus (Bosco-Lauth et al., 2018, 2016; Shah et al., 1966). To investigate its naturally-occurring host range, a user might begin by loading the data, subsetting to records about chikungunya virus, and examining the raw data:

~~~
### R Script:
### Note that in these examples we rely heavily on the R package family ‘tidyverse’ for ease of manipulating large data
> library(tidyverse); library(vroom)
> virion <- vroom(“Virion/virion.csv.gz“)
> virion %>%
+ filter(Virus == “chikungunya virus“) %>%
+ View()
~~~

There are nearly 2,000 records for chikungunya virus in the VIRION database that are called up by this function. If a user was interested in investigating the host range contained in those records, they might then call:

~~~
> virion %>%
+ filter(Virus == “chikungunya virus“) %>%
+ select(Host) %>%
+ n_distinct() # Reports number of unique hosts
[1] 27
> virion %>%
+ filter(Virus == “chikungunya virus“) %>%
+ select(HostOrder) %>%
+ distinct() # Reports the host orders
# A tibble: 4 × 1
HostOrder
<chr>
1 passeriformes
2 primates
3 chiroptera
4 rodentia
~~~

There are (at the time of writing) 27 unique hosts, including primates, bats, rodents, and - surprisingly - perching birds (passerines). Interested in following up on that finding, a user could simply call:

~~~
> virion %>%
+ filter(Virus == “chikungunya virus”, HostOrder == “passeriformes“) %>%
+ select(Host, Database, DetectionMethod)
# A tibble: 2 × 3
Host Database DetectionMethod
<chr> <chr> <chr>
1 passer luteus Shaw Isolation/Observation
2 passer luteus GLOBI Not specified
~~~

In most cases, viral isolation is taken as the most stringent possible evidence of host competence based on detection alone (Becker et al., 2020b), implying that the Sudan golden sparrow (*Passer luteus*) or species like it could be part of the virus’s sylvatic cycle in Africa. (This example also highlights another key point about the VIRION architecture: because some data sources index others – in this case, because the Shaw database is indexed in GLOBI – there will be some redundancy between records. While this is to be expected, it can also lead to metadata loss; for example, here, the GLOBI records do not distinguish viral isolation from PCR or serology.) Despite this interesting bird record, mammals seem a more likely sylvatic host on balance (though the distribution of research effort has also probably been uneven):

~~~
> virion %>%
+ filter(Virus == “chikungunya virus”,
+ DetectionMethod == “Isolation/Observation“) %>%
+ select(Host, HostOrder) %>%
+ group_by(HostOrder) %>%
+ summarize(nHost = n_distinct(Host)) %>% # Calculate unique hosts per order
+ arrange(-nHost)
# A tibble: 4 × 2
HostOrder nHost
<chr> <int>
1 chiroptera 5
2 primates 4
3 rodentia 2
4 passeriformes 1
~~~

While mosquito-borne alphaviruses like Mayaro virus are usually thought of as primate-reservoired, these results suggest that closer attention might be required with regard to the role that bats play in the circulation of chikungunya — an idea with broader support for other arboviruses (Fagre and Kading, 2019).

#### Example 2: Do fish host any zoonotic viruses?

Most zoonotic viruses are hosted by mammals, while a handful have bird reservoirs; only a small number can infect other vertebrate classes. Are there any truly zoonotic viruses that can also infect fish? To answer this question, we can first generate a list of viruses that are recorded as infecting humans:

~~~
> virion %>%
+ filter(Host == “homo sapiens“) %>%
+ select(Virus) %>%
+ distinct() %>%
+ pull(Virus) -> zoonoses
~~~

Next, we pull all records of those viruses in fish:

~~~
> fish <- c(“actinopteri”, “chondrichthyes”, “myxini”, “hyperoartia“)
> virion %>%
+ filter(Virus %in% zoonoses) %>%
+ filter(HostClass %in% fish) %>%
+ select(Virus) %>%
+ distinct() %>%
+ pull(Virus)
[1] “infectious spleen and kidney necrosis virus” “canine morbillivirus”
[3] “australian bat lyssavirus” “vesicular exanthema of swine virus”
[5] “norovirus sp.” “enterovirus sp.”
[7] “circoviridae sp.” “cress virus sp.”
[9] “simian immunodeficiency virus” NA
[11] “variola virus”
~~~

Four of these are unresolved at the species level (despite valid species-level identifiers in the NCBI taxonomy); these include classic food-borne pathogens found in sewage (enteroviruses and noroviruses) and poorly-characterized single-stranded DNA viruses (circoviruses and circular rep-encoding single-stranded, or CRESS, viruses). But the remaining six results are surprising, and each tells a slightly different story about how users should work with the types of data stored in VIRION.

1. **Variola virus:** Variola virus - more commonly known as smallpox - is a now-eradicated human infectious disease. Although some animals have been experimentally infected, no evidence suggests that smallpox has any unknown zoonotic reservoirs or notable natural animal hosts to consider; why is this virus being reported from fish? We can start by tracing the record to the source:

~~~
> virion %>%
+ filter(Virus == “variola virus”,
+ HostClass %in% fish) %>%
+ select(Host, HostGenus, HostFamily, HostOriginal, Virus, Database) %>%
+ unique()
# A tibble: 1 × 6
Host HostGenus HostFamily HostOriginal Virus Database
<chr> <chr> <chr> <chr> <chr> <chr>
1 NA variola serranidae Variola variola virus GLOBI
~~~

Here, GLOBI has recorded an erroneous self-interaction between this virus and itself as a “host”, likely due to an error in the processing of an automated scraping algorithm: because the record places “Variola” in the Host field, our taxonomic pipeline assigns the record to the only vertebrate host genus for *Variola*, the lyretails (Serranidae).
2. **Australian bat lyssavirus:** Here, we can again start by checking the source of the records that place a bat virus in a fish host:

~~~
> virion %>%
+ filter(Virus == “australian bat lyssavirus”,
+ HostClass %in% fish) %>%
+ select(Host, DetectionMethod, Database, ReferenceText) %>%
+ unique()
# A tibble: 2 × 4
Host DetectionMethod Database ReferenceText
<chr> <chr> <chr> <chr>
1 epalzeorhynchos kalopterus PCR/Sequencing EID2 NCBI Nucleotide
2 epalzeorhynchos kalopterus PCR/Sequencing GenBank NA
> virion %>%
+ filter(Virus == “australian bat lyssavirus”,
+ HostClass %in% fish) %>%
+ pull(NCBIAccession)
[1] “14162205” “14162207” “14162209” “14162211” “AF369369” “AF369370” “AF369371” “AF369372”
~~~

GenBank appears to be the original source for this record. There are many pitfalls that users can encounter when submitting data to sources like GenBank. What’s going on with these samples?

~~~
> nuc <- rentrez::entrez_summary(db = “nuccore”, id = “14162205“)
> nuc$subtype
[1] “isolate|host”
> nuc$subname
[1] “V474.ABL|Flying fox”
~~~

This is the result of an unfortunate ambiguity in the original submission. This host common name is a synonym in the NCBI taxonomy with the fish host *Epalzeorhynchos kalopterus,* which is in fact called the flying fox (TaxID: 699555); this can easily be confirmed with taxize::classification(“Flying fox”, db = “ncbi“). The original data should instead refer to the flying fox bat by its scientific name (*Pteropus* sp.).
3. **Simian immunodeficiency virus:** This record comes from a similar problem with user-submitted ambiguous data. Here, a user has submitted the host name “Rita”, which has unfortunately matched to the fish genus *Rita* (TaxID: 337744):

~~~
> virion %>%
+ filter(Virus == “simian immunodeficiency virus”,
+ HostClass %in% fish) %>%
+ select(Host, DetectionMethod, Database, NCBIAccession) %>%
+ unique()
# A tibble: 2 × 4
Host DetectionMethod Database NCBIAccession
<chr> <chr> <chr> <chr>
1 NA PCR/Sequencing GenBank AY932804
2 NA PCR/Sequencing GenBank AY932808
> rentrez::entrez_summary(id = “AY932804”, db = “nuccore“)$subtype
[1] “isolate|host|gb_acronym|country|isolation_source”
> rentrez::entrez_summary(id = “AY932804”, db = “nuccore“)$subname
[1] “SIVsmTAI-23|Rita (sooty mangabey)|SIV|Cote d’Ivoire: Tai National
Park|stool from wild-living sooty mangabey”
~~~

This is a bit of a strange error, given that *Rita* is not an alternate genus or outdated genus name for this species (*Cercocebus atys*). Who, or what, is Rita? A quick inspection of a related publication(Santiago et al., 2006) reveals that Rita was the name assigned to an adult female monkey from a monitored population of sooty mangabeys in Côte d’Ivoire.
4. **Infectious spleen and kidney necrosis virus:** This record has a slightly different kind of problem: this is a well-studied fish virus, but not one ever known to cause human disease. Why is it included on our list of zoonotic viruses? This also appears to be an erroneous GenBank accession, which – as with the previous example – has been propagated elsewhere:

~~~
> virion %>%
+ filter(Virus == “infectious spleen and kidney necrosis virus”,
+ Host == “homo sapiens“) %>%
+ select(Host, DetectionMethod, Database, ReferenceText) %>%
+ unique()
# A tibble: 2 × 4
Host DetectionMethod Database ReferenceText
<chr> <chr> <chr> <chr>
1 homo sapiens PCR/Sequencing EID2 NCBI Nucleotide
2 homo sapiens PCR/Sequencing GenBank NA
~~~

The host assigned to the relevant accessions (e.g., HQ317457.1) is *Homo sapiens*, for reasons that cannot be easily reconstructed; the source publication (Fu et al., 2011) only records samples from fish species. This may be the result of an automated pipeline error or something similar, but is difficult to resolve, and would be a case to directly contact the authors of these samples with further queries.
5. **Canine morbillivirus:** These records also highlight the challenges of user-submitted GenBank data, but are less identifiably the result of a user-end error.

~~~
> virion %>%
+ filter(Virus == “canine morbillivirus”,
+ HostClass %in% fish) %>%
+ select(Host, DetectionMethod, Database, ReferenceText) %>%
+ unique()
# A tibble: 4 × 4
Host DetectionMethod Database ReferenceText
<chr> <chr> <chr> <chr>
1 coregonus autumnalis PCR/Sequencing EID2 NCBI Nucleotide
2 cottocomephorus grewingki PCR/Sequencing GenBank NA
3 coregonus migratorius PCR/Sequencing GenBank NA
4 coregonus autumnalis Not specified GLOBI NA
> virion %>%
+ filter(Virus == “canine morbillivirus”,
+ HostClass %in% fish) %>%
+ pull(NCBIAccession) %>%
+ unique()
[1] “209961611” “209961613” “EU787998” “EU787999” “EU788000” “EU788001” NA
~~~

A quick check will confirm all of these samples refer to a single study (Butina et al., 2010) on canine distemper in the Lake Baikal seal (*Phoca sibirica*). Though the study makes no note of samples from these species, the samples seem unlikely to be user-end errors. For example, user-submitted metadata from accession EU787998 not only records the host as “Cottocomephorus grewingki (Dybowski, 1874)” but also name the isolate as “Cottoc9-06”; these details suggest it is unlikely these entries were made in error, but without additional information from the peer-reviewed study or the authors, it is difficult to make much further inference.
6. **Vesicular exanthema of swine virus:** Of the six fish viruses we examined, this is perhaps the only one that seems strongly supported at face value; peer-reviewed studies have described the isolation of the virus from both the opaleye perch (Smith et al., 1980) and humans (Smith et al., 1998):

~~~
> virion %>%
+ filter(Virus == “vesicular exanthema of swine virus”,
+ HostClass %in% fish) %>%
+ select(Host, DetectionMethod, Database, ReferenceText) %>%
+ unique()
# A tibble: 1 × 4
Host DetectionMethod Database ReferenceText
<chr> <chr> <chr> <chr>
1 girella nigricans Isolation/Observation Shaw Smith AW, 1980, Science, 209
> virion %>%
+ filter(Virus == “vesicular exanthema of swine virus”,
+ Host==“homo sapiens“) %>%
+ select(Host, DetectionMethod, Database, ReferenceText) %>%
+ unique()
# A tibble: 8 × 4
Host DetectionMethod Database ReferenceText
<chr> <chr> <chr> <chr>
1 homo sapiens Not specified EID2 Neill JD, 1995, J Virol, 69
2 homo sapiens Not specified EID2 Lazo A, 2002, Vox Sang, 83
3 homo sapiens Not specified EID2 Chen R, 2006, Proc Natl Acad Sci U S A, 103
4 homo sapiens PCR/Sequencing EID2 NCBI Nucleotide
5 homo sapiens Isolation/Observation Shaw Smith AW, 1998, Clin Infect Dis, 26
6 homo sapiens Isolation/Observation Shaw ViPR
7 homo sapiens PCR/Sequencing GenBank NA
8 homo sapiens Not specified GLOBI NA
~~~

Vesicular exanthema of swine virus (VESV) is an unusual virus. The species *sensu lato* includes a number of closely related caliciviruses found in cattle, marine mammals, and even reptiles. Work from the 1990s reports that the virus can cause disease in humans (Smith et al., 1998), who appear to sometimes acquire the infection through exposure to sewage-contaminated shellfish; however, the Wikipedia for VESV currently states that the virus is “not transmissible to humans,” highlighting the value of VIRION as a resource to point to these older records. Given how many viruses are classified under VESV at the species rank, it may be that the lineages that infect fish and humans are quite genetically distinct, and may eventually be reclassified as separate species; currently, the ICTV recognizes VESV and feline calicivirus as the two valid species in the genus (Caliciviridae: *Vesivirus*), as well as a handful of other unclassified viruses. For now, among the possible candidates for a “fish-borne zoonotic virus” in VIRION, VESV is certainly the strongest contender.

### Anticipated Future Improvements

We aim to maintain and expand VIRION to keep pace with the rapidly emerging field of viral ecology. The existing dataset has notable opportunities for future metadata improvement; for example, efforts to manually or programmatically revisit source publications may help flag experimental infections, and better differentiate host-virus associations that are only biologically possible from those that already occur in nature. Similarly, future iterations of the dataset might incorporate new kinds of taxonomic designations, particularly if it becomes possible to efficiently implement OTU-based designations from sequence data at scale. In all of these cases, researchers can expect minor announcements about changes in workflow, scope, and format to be made via GitHub release notes, while more sizable updates could also include a blog post on the website of the Verena Consortium (viralemergence.org) or, in the case of major expansions, an additional scientific publication.

Keeping VIRION useful to researchers will also require ongoing efforts to better integrate existing sources (e.g., the continuously-updated EID2 database, if the web interface is updated to allow full downloads) and new sources developed in the future. VIRION is intended to be the flagship database at the heart of a broader open data ecosystem developed by the Verena Consortium, and will eventually be integrated with other existing databases (e.g., the bat betacoronavirus host database: viralemergence.org/betacov) and other datasets yet-to-be-developed. The current scope of VIRION could also lay the foundation to incorporate other data sources beyond vertebrate viruses; for example, we have developed the ‘insectDisease’ R package to access the Ecological Database of the World’s Insect Pathogens (github.com/viralemergence/insectDisease), but some of these data could be easily incorporated into a future iteration of VIRION that includes all animals. Similarly, a specialized extension of VIRION may be useful to describe vector-arbovirus relationships, which have been catalogued in a handful of publications (e.g., (Evans et al., 2017)) and may benefit from tailored data architectures. There may even come a point where researchers or teams (e.g., broadly-coordinated efforts to discover wildlife viruses) may want to deposit association data as they are gathered directly into VIRION, particularly if these data are used to power model-guided sampling strategies that could accelerate viral discovery (Becker et al., 2020a). All of these may be future priorities for VIRION development.

## Acknowledgements

This work was supported by funding to the Viral Emergence Research Initiative (VERENA) consortium, including NSF BII 2021909. Part of this contribution was enabled by support provided by Calcul Québec (www.calculquebec.ca) and Compute Canada (www.computecanada.ca). Ryan Connor is supported by the National Library of Medicine, National Center for Biotechnology Information at the National Institutes of Health. The authors thank the National Center for Biotechnology Information for extensive informatic and research support during the design process, and all of the thousands of scientists who have contributed the data that form the backbone of the VIRION project.

## Conflicts of interest

The authors declare no conflicts of interest. This material should not be interpreted as representing the viewpoint of the U.S. Department of Health and Human Services, the National Institutes of Health, National Library of Medicine, National Center for Biotechnology Information, Center for Information Technology.

## Data and code availability

The VIRION database, and the code used to generate it, are available from the Verena Consortium’s GitHub account at github.com/viralemergence/virion. Additional code used to produce the CLOVER dataset is available at github.com/viralemergence/clover.

## Author contributions

CJC, GFA, and TP conceived the study. CJC and RG led data architecture development, and CJC, RG, GFA, RPC, TAD, MJF, and TP contributed code. CJC, RLM, MJF, and RG generated visualizations. All authors contributed to beta-testing and to the manuscript.

## Notes

http://www.github.com/viralemergence/virion

